# A clinically annotated post-mortem approach to study multi-organ somatic mutational clonality in histologically healthy tissues

**DOI:** 10.1101/2022.02.27.482170

**Authors:** Tom Luijts, Kerryn Elliott, Joachim T. Siaw, Joris Van de Velde, Elien Beyls, Arne Claeys, Tim Lammens, Erik Larsson, Wouter Willaert, Anne Vral, Jimmy Van den Eynden

## Abstract

Recent research on histologically healthy human tissues identified omnipresent mutational microclones, driven by somatic mutations known to be responsible for carcinogenesis (e.g., in *TP53* or *NOTCH1*). These new insights are fundamentally changing current tumour evolution models, with broad oncological implications. Most studies are based on surgical remnant tissues, which are not available for many organs and rarely in a pan-organ setting (multiple organs from the same individual). Here, we describe an approach based on clinically annotated post-mortem tissues, derived from whole-body donors that are routinely used for educational purposes at human anatomy units. We validated this post-mortem approach using UV-exposed and unexposed epidermal skin tissues and confirm the presence of positively selected *NOTCH1/2-, TP53*- and *FAT1*-driven clones. No selection signals were detected in a set of immune genes or housekeeping genes. Additionally, we provide the first evidence for smoking-induced clonal changes in oral epithelia, likely underlying the origin of head and neck carcinogenesis. In conclusion, the whole-body donor-based approach provides a nearly unlimited healthy tissue resource to study mutational clonality and gain fundamental mutagenic insights in the presumed earliest stages of tumour evolution.

## Introduction

Spontaneously occurring somatic mutations accumulate in the genome of dividing cells during ageing. Most of these mutations are harmless, but occasionally, they lead to a cellular fitness advantage, resulting in positive selection and clonal expansion. The progressive accumulation of these driver mutations can lead to malignant tumour formation, often many decades after the occurrence of the first driver event^1,2^. International efforts like The Cancer Genome Atlas (TCGA) and International Cancer Genome Consortium (ICGC) led to the identification of these genomic alterations in most primary tumour types. In contrast, less is known about the earliest initiating events, which are expected to occur in normal cells.

Mutational clonality in histologically healthy tissues was first shown for *TP53* in the UV-exposed skin using immunohistochemical techniques^3^. However, it took more than 20 years and the development of advanced Next-Generation Sequencing (NGS) techniques before these findings could be extended to other cancer genes and tissue types. In their pioneer study, *Martincorena et al*. used a deep (500x) targeted sequencing approach on 74 genes in 4 subjects and demonstrated that 18-32% of all UV-exposed epidermal skin cells are (micro-) clones that are driven by point mutations in cancer genes commonly found in squamous cell carcinoma (SCC) skin cancer, such as *TP53* and *NOTCH1*^4^. These findings were later confirmed in healthy oesophageal tissues, where age- and smoking-dependent mutational clonality was demonstrated in genes known to be involved in oesophageal squamous cell carcinoma (ESCC)^5,6^. Recently, similar DNA-based studies were performed on non-tumoral colorectal^7,8^, uterine^9,10^, liver^11,12^, lung^13^ and urothelial tissues^14,15^ and an indirect RNA-based approach found clonal expansion in several healthy tissue types^16^. The high frequency and large size of these clones is striking, as illustrated by *NOTCH1*-driven clones in ESCC. These results suggest that some genes favour the initial clonal expansion in normal tissues without the risk of further evolution into cancer^17^. Furthermore, clones with different carcinogenic potential might compete for space, especially within aging tissues^18,19^. These new carcinogenic insights also have major translational relevance. They could even imply that therapeutically targeting genes like *NOTCH1* might increase the risk of malignant transformation by tackling benign rather than malignant clones, and opposite therapeutic approaches should be considered. Additionally, the implications for diagnostic strategies based on e.g., circulating tumour cells or cell-free DNA are to be determined. There is also a non-oncological significance as mutational clonality has been suggested to contribute to aging and other human diseases^12,20^. In this regard it has been shown that mutational clonality in blood cells has a negative, cancer-independent impact on lifespan, likely related to an increased incidence of cardiovascular diseases^21^.

An important limitation for future studies is the lack of clinically annotated normal tissue availability. Current studies are mostly based on remnant tissues, obtained from surgical procedures. This approach is not feasible for all tissues and a drawback is the lack of comparability between different organs when samples are not obtained from the same patient. In this study, we describe an approach based on post-mortem tissues obtained from whole-body donors. Using deep targeted sequencing, we demonstrate UV-induced genomic alterations and confirm positive selection of somatic mutations in *TP53* and *NOTCH1* in epidermal skin tissues. Our study also provides the first evidence for smoking-induced somatic alterations in oral tissues. The newly developed approach potentiates future studies focussing on multi-organ genomic effects of cancer risk factors and mutagens (e.g., smoking, radiotherapy, chemotherapy), providing a valuable tool to study human carcinogenesis and other age-related diseases.

## Results

### A whole-body donor-based approach to study mutational clonality

On average 101 bodies are voluntarily donated to the Ghent University Anatomy and Embryology Unit each year (data 2016-2021; range 82-133). The median age of these whole-body donors is 81 years (range 36 – 106 years; Fig. 1a) and the majority (53%) deceased between 12 and 36 hours prior to arrival (post-mortem interval; PMI), with 92% of all donors having a PMI below 72 hours (Fig. 1b). They are primarily used for medical and surgical training purposes^22,23^, but also provide a rich normal tissue resource to study clonal changes in different organs of the human body.

**Fig. 1.**
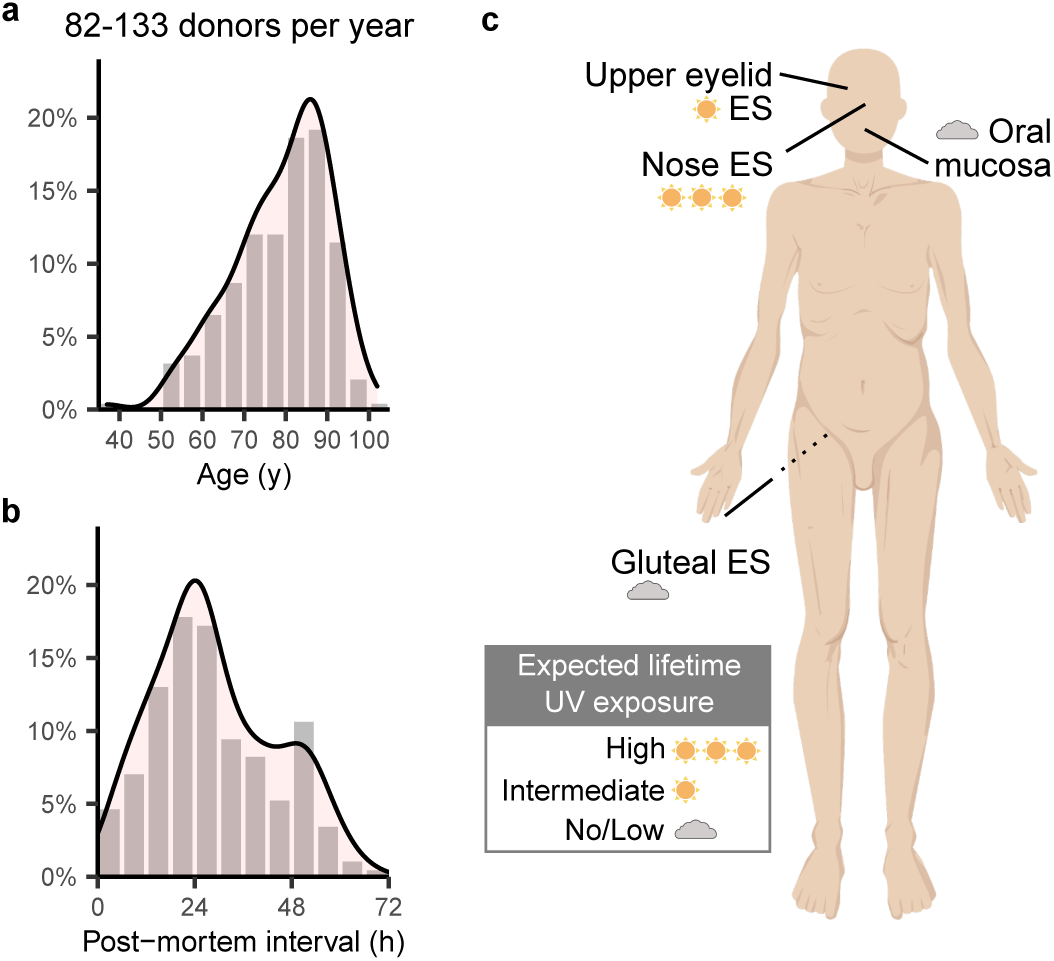
Detecting mutational clonality in post-mortem epithelial tissues derived from whole-body donors. **(a-b)** Histograms showing the distribution of **(a)** age and **(b)** post-mortem interval in the complete population of whole-body donors at the UGent Anatomy and Embryology unit (data 2016-2021). **(c)** Skin and oral tissues were sampled from 4 different locations as indicated.

To demonstrate the feasibility of such a post-mortem tissue-based mutational clonality detection platform, we developed a 4-step methodology (Fig. S1a). In a first step, skin and oral tissues were sampled and relevant clinical information (i.e., age, gender, smoking status, oncological history) as well as the PMI was recorded by the responsible physician. The choice for skin tissues for initial evaluation purposes was motivated by their easy accessibility, high expected UV-induced mutation rates (with negative controls from unexposed regions) and knowledge of the expected (positive control) results from earlier studies^4^. Secondly, 5 mm (diameter) punch biopsies were taken (surface area 19.63 mm^2^), and the epithelial layer was enzymatically isolated. In a third step DNA was extracted, DNA concentrations and integrity were determined, and samples were sequenced. Importantly, we did not observe any PMI-dependent DNA degradation, as evaluated using gel electrophoresis (eyelid samples in 10 subjects with PMI between 6h and 83h evaluated, Fig. S1b). In a fourth step, somatic mutations were called, followed by an analysis of clone sizes, mutational signatures, and positive selection signals.

We aimed to validate our approach using 2 largely orthogonal approaches: 1) by identifying positive selection of somatic mutations in cancer genes and 2) by demonstrating UV-specific alterations in UV-exposed skin. To achieve this, we sampled skin and oral tissues from 2 different donors: a 94-year-old, male, non-smoking donor (subject PM01, PMI 33h) and a 90-year-old, male donor (subject PM02, PMI 38h) with a smoking history (1 package a day between the age of 18 and 38, as derived from the medical record). To explore the effects of different expected lifetime UV exposure, skin samples were taken at 3 locations: bridge of the nose (high UV exposure), upper eyelid (intermediate UV exposure) and gluteal region (no/low UV exposure). Additional non-UV-exposed oral mucosa samples were taken from the inner buccal region (Fig. 1c).

### Somatic mutations in known driver genes are detectable in post-mortem tissues

Deep (1000x) targeted sequenced was performed on 153 genes. Apart from 76 earlier defined skin cancer genes, this gene panel also contained 1) 25 driver genes that have been associated to skin or head and neck cancer, 2) 20 housekeeping control genes and 3) 32 genes that are putatively involved in immune evasion (table S1). After alignment, somatic mutations were called using *Shearwater ML*, an algorithm that is optimized for the detection of low frequency variants from deep targeted sequencing data. We retrieved 910 somatic mutations with an average variant allele frequency (VAF) of 0.011 and ranging between 0.0023 and 0.11 (Table S1).

The 5 most frequently mutated genes were *NOTCH1* (6.6 average mutations per sample; mps), *TP53* (3.1 mps), *MUC17* (2.5 mps), *FAT1* (2.2 mps) and *APOB* (2.1 mps). Most samples contained multiple mutations in these genes, and remarkably, *NOTCH1* mutations were identified in all samples (between 1 and 16 different *NOTCH1* mutations per sample). Overall, we identified somatic mutations in 119 different genes, including *NOTCH2* (1.6 mps, 6^th^ most frequently mutated gene) and *NOTCH3* (0.82 mps), genes that have been previously identified as clonal drivers in healthy skin (Fig. 2)

**Fig. 2.**
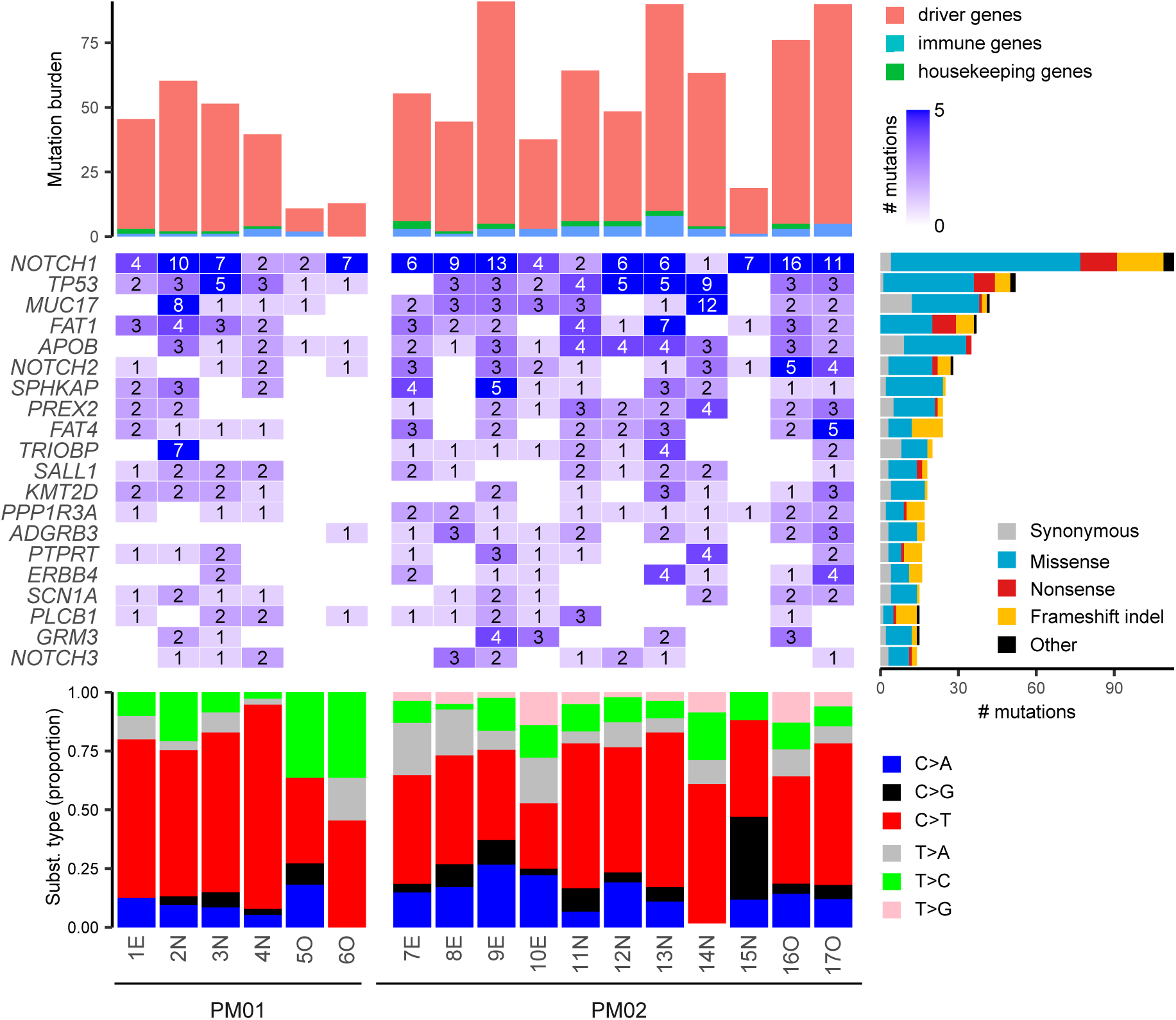
Characterization of somatic mutations in epidermal skin and oral epithelia. Somatic mutations were called from deep (1000x) targeted sequencing. The 20 most frequently mutated genes with indication of the number of identified mutations per sample are shown in the middle plot. Genes are ranked from top to bottom following mutation frequency, as indicated by the right bar. Upper plot shows the mutation burden, stacked according to the type of gene (driver gene, immune gene or housekeeping gene, as indicated). Bottom plot shows the distribution of the 6 main substitution types in each sample. Sample and patient pseudo-identifiers indicated on the bottom. E, epidermal eyelid sample; N, epidermal nose sample; O, oral sample.

Most of these mutations were observed in the 101 driver genes (93%, 844 mutations). The majority of the remaining mutations occurred in the 32 immune genes (46 mutations, 5%), while 20 mutations (2%) were observed in the 20 housekeeping control genes (Fig. S2, table S1).

### Clonal alterations in epidermal skin and oral mucosa are primarily driven by *NOTCH1* and *TP53*

The large difference between the number of mutations observed in driver versus housekeeping genes suggests positive selection acting on the former. To confirm this, we first focused on the global ratio of nonsynonymous over synonymous mutations. In skin samples, this ratio was higher than the expected ratio, as derived from a random mutation background model (3.68 versus 2.12 respectively; *P* = 3.7e-08). This higher observed to expected mutation ratio (i.e., global dN/dS = 1.73) suggests that 42% (0.73/1.73) of the identified nonsynonymous mutations have been subjected to positive selection forces (Fig. 3a). Significant selection signals (i.e., dN/dS values above 1) were solely observed in the group of driver genes (dN/dS = 1.78, *P* = 2.4e-08), and not in the immune genes (dN/dS = 0.81, *P* = 0.72) nor the housekeeping genes (dN/dS = 0.65, *P* = 0.81; Fig. 3b). Additionally, signals were stronger for nonsense mutations (dNons/dS = 3.04, *P* = 3.9e-09) than for missense mutations (dMiss/dS = 1.80, *P* = 6.3-09) and were also present in the oral samples (dN/dS = 2.13, *P* = 1.3e-04; Fig. 3a-b, Table S1). When focusing on single genes in skin, we observed significant selection signals (at 10% FDR) in *NOTCH1, TP53* and *FAT1* (Fig. 3a, Fig. S3).

**Fig. 3.**
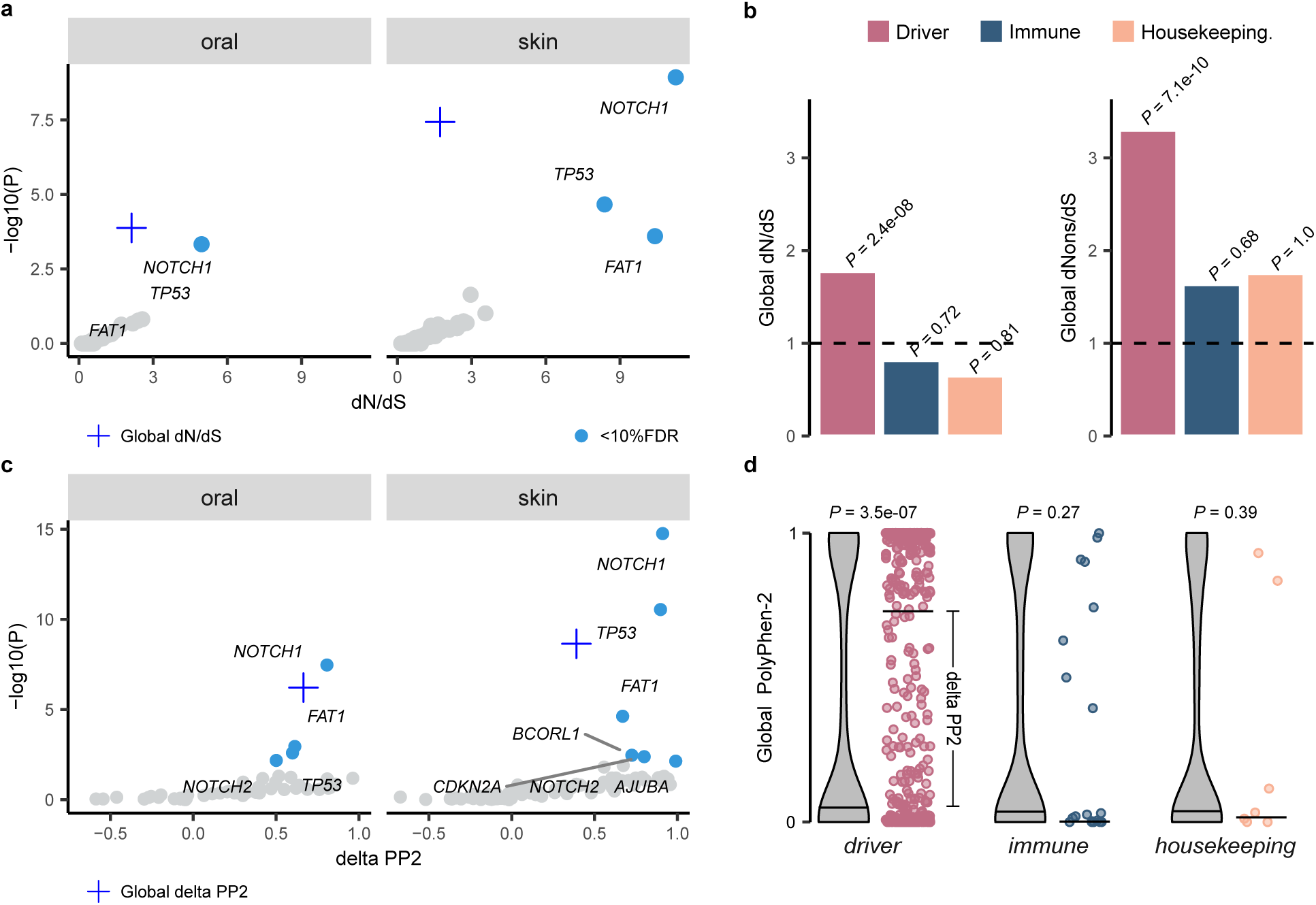
Positive selection signals in driver mutations in oral and skin epithelial tissues. **(a-b)** Analysis of the number of substitutions. dN/dS and dNons/dS values calculated by normalization to the expected number of sites and *P* values calculated using a one-sided binomial test (*Methods*). **(a)** Scatter plot with dN/dS values on the x-axes and (log10-transformed) *P* values on the y-axes. Genes containing significant dN/dS values (at 10% FDR) indicated in blue. Global dN/dS value indicated by plus-sign. **(b)** Barplot showing global dN/dS or global dNons/dS for driver, immune and housekeeping gene sets as indicated. **(c-d)** Analysis of the PolyPhen-2 (PP2) functional impact scores. For each gene, PP2 scores were compared with their expected value as simulated in the absence of any selection pressure (*Methods*). *P* values calculated using a one-sided Wilcoxon rank sum test. **(c)** Scatter plot with skin delta PP2 (difference between median observed and expected values, as illustrated in panel d) on the x-axes and *P* values on the y-axis. Genes with significant differences (at 10% FDR) indicated in blue. **(d)** Plots showing expected (violin plot) and observed (scatter plots) PP2 values for skin driver, immune and housekeeping gene sets as indicated. Median values indicated by horizontal lines.

Because selection pressures are mainly expected on nonsynonymous mutations with high mutational impact, we also compared PolyPhen-2 (PP2) mutational impact scores between observed and expected mutations. Higher PP2 scores were observed for our set of somatic mutations (median PP2 = 0.50) as compared to the expected scores (median PP2 = 0.11; *P* = 2.3e-09; Fig. 3c). Like the dN/dS approach, this difference was only observed for driver genes (*P* = 3.5e-07) and not for the other gene sets (Fig. 3d). We confirmed positive selection signals (at 10% FDR) in skin tissues in *NOTCH1, TP53* and *FAT1* and, additionally, also found significant signals in *NOTCH2, CDKN2A, BCORL1* and *AJUBA* (Fig. 3c, Fig. S3). In oral tissues, positive selection was detected in *NOTCH1, FAT1, TP53* and *NOTCH2* (Fig. 3c).

Our results confirm earlier reports on somatic driver mutation clonality in healthy skin. Based on the variant allele frequency and biopsy size, we imputed the clone sizes for the genes that were identified to be under positive selection and found 78 clones per cm^2^ skin (35 *NOTCH1*, 16 *TP53*, 15 *FAT*, 6 *NOTCH2*, 2 *BCORL1*, 2 *CDKN2A* and 2 *AJUBA* clones; Fig. 4a, b). *BCORL1*-driven clones had the largest size (median 0.49 mm^2^) while the smallest clone sizes were found for *FAT1* (0.23 mm^2^; Fig. 4a, c). These clone sizes were higher than described previously^4^, although the total percentage of skin occupied by the most abundant clones was largely comparable between both studies, with 15.2% of skin cells estimated to contain *NOTCH1* mutations, 5.7% *TP53* mutations, 4.7% *FAT1* mutations and 2.1% *NOTCH2* mutations (Fig. 4d). In the oral mucosa, similar clonal densities were observed for the donor with the smoking history, but not for the non-smoking donor, where clones were relatively scarse (Fig. S4).

**Fig. 4.**
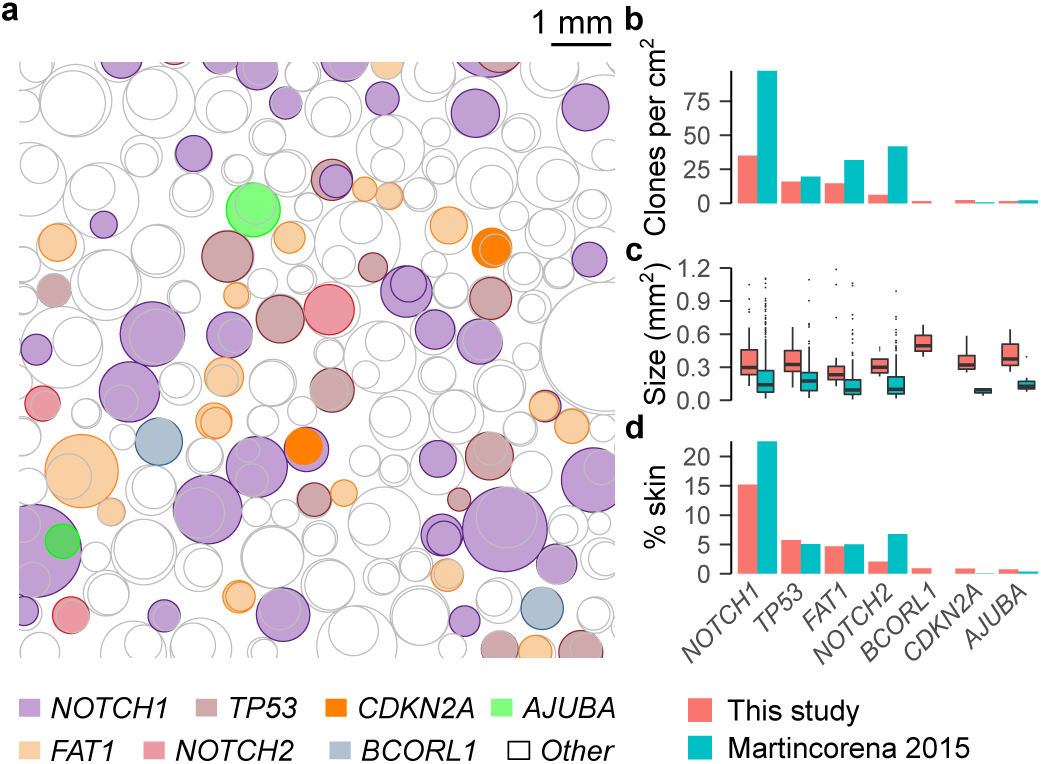
Clonal alterations in healthy UV-exposed skin. **(a)** Visualization of the estimated number and size of somatic mutation-driven clones in 1 cm^2^ healthy skin. Data from PM01 and PM02 pooled. Clones driven by genes for which positive selection signals were found in this study are coloured as indicated in bottom legend. Other clones are uncoloured. Clones were positioned randomly with clone sizes and frequency based on data from this study. **(b-d)** Comparison of **(b)** clone frequency (number per cm^2^), **(c)** clone size and **(d)** estimated percentage of skin occupied by each clone between this study and the study from *Martincorena* 2015 for genes for which positive selection signals were found.

### UV-specific genomic alterations in epidermal skin correlate to the expected amount of lifetime UV exposure

As an additional validation of our approach, we aimed to detect UV-induced genomic changes in post-mortem tissues derived from whole-body donors. The substitutions in the 12 skin samples were predominated by C>T mutations (56%; Fig. 2, Fig. 5a). Contrary to the other 5 substitution types and as expected for UV-induced somatic mutations, these C>T mutations mainly occurred in a dipyrimidine context (88%) and were 24% more prevalent in the coding strand than the template strand (*P* = 0.054, exact Poisson test), suggesting transcriptional strand bias (Fig. 5a). This mutational pattern is consistent with previous UV exposure, which was further confirmed by the strong predominance (82.4%) of coding strand-specific CC>TT dinucleotide variants (DNVs; *P* = 3.4e-10; Fig. 5b) and similarity of the samples’ 96 trinucleotide substitution type signature (i.e., substitution type and adjacent base pairs) with the well-known UV-associated single base substitution signature (SBS) 7 (Fig. 5c). This signature was found in all skin samples (mean contribution 39.3% per sample) and was the most prevalent signature in 10/12 skin samples. Interestingly, the smoking-related signature 4 was retrieved in 7/11 samples from subject PM02, which was known to have a smoking history (Fig. 5c).

**Fig. 5.**
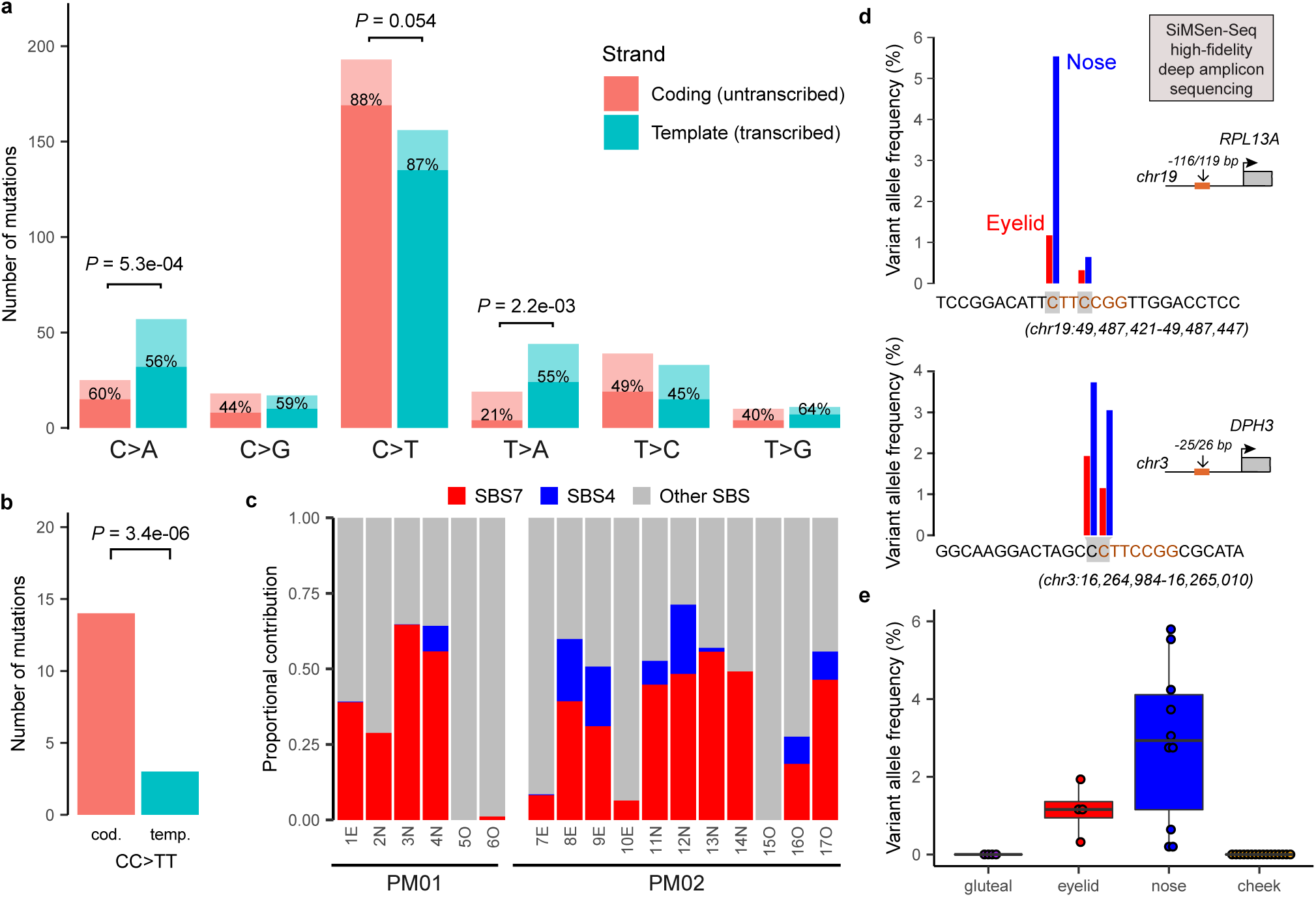
Detection of UV-induced somatic mutations in post-mortem epithelial tissues. **(a-c)** Deep (1000x) targeted sequencing results. **(a-b)** Bar plots representing the total number of single nucleotide variants (**a**) or dinucleotide variant (DNVs; **b**), detected in epidermal skin (12 samples). Substitutions are stratified according to the occurrence of the pyrimidine on the coding (untranscribed) or template (transcribed) strand as indicated. Transcriptional strand bias tested using Poisson exact test. Bars are coloured by strand as indicated and following the number of substitutions occurring in a dipyrimidine (dark, % indicated) or other sequence (light) context. **(c)** Proportion of known mutational signatures retrieved by analysing the cosine similarity between the distribution of 96 trinucleotide substitution types and 30 known COSMIC mutational signatures. Single base substitution signatures (SBS) 4 and 7 coloured as indicated. **(d-e)** SiMSen sequencing of well-known UV hotspot mutations in *RPL13A* and *DPH3* promoters. (**d**) Data from PM02 for illustration, 2 hot spots each as indicated by grey shaded nucleotides, sequence indicated in orange. (**e**) Boxplot summarizes results for both study subjects, all hotspots and sample locations as indicated. Boxplot indicates median values and lower/upper quartiles with whiskers extending to 1.5x the interquartile range. Dots show individual data points.

To exclude that this putative UV signal was biased by the rather limited genomic coverage of our gene panel and/or positive selection processes, we extended our analysis to known UV hotspots in the promoter regions of *RPL13A* and *DPH3*. Here, C>T mutations occurring in a **<u>CC</u>**TT**<u>C</u>**CGG sequence context have previously been associated with the amount of cumulative UV-exposure due to ETS transcription factor binding^24,25^. We sequenced these regions using an error-correcting amplicon sequencing protocol (SiMSen-Seq) and detected UV-specific mutations in the UV-exposed eyelid and nose samples, but not in the non-exposed gluteal or oral samples (Fig. 5d, Table S1). Further, the variant allele frequency (VAF) was significantly higher in nose samples than eyelid samples, in line with the higher expected lifetime UV exposure of the former (Fig. 5e). These results confirm that low frequency UV-induced mutations can be accurately detected in post-mortem skin tissues.

## Discussion

Omnipresent epithelial microclones have been identified in different organs^26^. These clones are driven by somatic mutations known to be involved in human carcinogenesis. New insights are changing our current understanding of (early) tumour evolution and have putative implications for cancer diagnostics and treatment, but further research is hampered by the rather limited availability of clinically annotated normal tissues. In this study, we have demonstrated that post-mortem tissues, derived from whole-body donors, could provide a nearly unlimited human tissue resource for future studies.

We validated our methodology on epidermal skin, a tissue where mutational clonality has been well-characterized and UV light is known to be an important mutagen. Firstly, we confirmed earlier findings on driver gene clonality. We found positive selection signals in *NOTCH1-2, TP53* and *FAT1*, demonstrating that these cancer genes are responsible for clonal alterations in healthy skin. While the estimated proportion of healthy skin cells containing these mutations (e.g., 15% *NOTCH1* mutations and 5% *TP53* mutations) was remarkably similar to previous reports based on either targeted sequencing approaches^4^ or immunohistochemistry^3^, the inferred clonal sizes and frequencies (number per cm^2^) were respectively higher and lower than reported previously. This is likely related to a lower sensitivity to detect small clones due to the larger punch biopsy sizes that were used in this study (5 mm diameter versus 1-2 mm in previous reports^4^). Indeed, assuming a minimal VAF detection threshold of 0.5% implies detectable clones of 19.6 mm^2^, 3.1 mm^2^ and 0.79 mm^2^ for 5 mm, 2 mm, and 1 mm punch biopsies, respectively. The theoretical advantage of larger punch biopsy sizes is the higher sensitivity to detect lowly frequent large clones. This could explain the additional identification of positive selection signals in *CDKN2A, BCORL1* and *AJUBA. CDKN2A* is a well-known tumour suppressor gene in several cancer types, including SCC^27^. *AJUBA* is a regulator of epidermal homeostasis and has been implicated in SCC as well^27,28^. Less is known about *BCORL1* (BCL6 corepressor like 1). Somatic mutations in this transcriptional corepressor have been identified in haematological malignancies^29^ and, interestingly, have been reported to be associated with resistance to *BRAF* inhibition treatment in melanoma^30^.

As a second validation approach, we confirmed UV-specificity of the identified genomic alterations in healthy skin. As expected, UV-exposed skin samples were characterized by a high amount of C>T single nucleotide mutations that occurred in pyrimidine dimers and were more frequent on the coding than the template strand. This transcriptional strand bias was most pronounced in the abundant CC>TT dinucleotide substitutions in skin. Further sequencing of 2 promoter regions that are known to be specifically altered by UV mutagenesis provided final confirmation.

Genomic immune evasion mechanisms (e.g., *B2M* mutations, *HLA* deletions) are relatively frequent in primary tumors^31,32^ and indirect evidence suggests their occurrence early during tumour evolution^33^, but this has never been shown directly. Therefore, we included a panel of genes in which somatic mutations have been suggested to result in immune evasion. We identified 46 mutations in these genes, but no selection signals were found. Larger studies are required to determine whether immune surveillance and evasion have any influence on epidermal skin microclonality.

One of the donors we sequenced was a former smoker, information that was derived from the medical record. Interestingly, we identified the cigarette smoke related SBS4 in most samples from this patient. Remarkably, this mutational signature was not restricted to the oral mucosa (where it is expected) but was also retrieved in facial skin samples. To our knowledge, these smoking-induced somatic mutations in healthy skin have not been identified previously. They could be related to the higher squamous skin cancer risk reported in smokers^34^, although no final conclusions are possible at this stage due to 1) the limited sample size of our study and 2) the rather limited genomic coverage of the gene panel with relatively large numbers of substitutions demonstrated to be under positive selection. Nevertheless, these findings clearly demonstrate the potential of a clinically annotated whole-body based approach to study somatic mutations in healthy skin.

Mutational clonality has been studied in post-mortem tissues derived from autopsy subjects^35^. However, autopsy approaches imply little or no control over the inclusion of subjects of interest. In contrast, the use of whole-body donors for future research on mutational clonality in healthy tissues offers great potential, particularly towards studies that aim to determine how human disease is directed by these clonal changes^12^ or how (iatrogenic) mutagens such as chemotherapy or radiotherapy influence driver mutational clonality in multiple organs. Our approach provides the opportunity to include subjects of interest by screening medical records upon donor arrival. Given the relatively high prevalence of oncological or cardiovascular diseases at the time of death, the inclusion of relatively large study populations is very realistic, especially when donor data from different human anatomy departments are merged as part of larger whole-body biobanking efforts.

## Methods

### Study subjects and ethics

Ten whole-body donors were included in this study between January 2020 and November 2021. Relevant clinical information (smoking status, oncological history, including chemo- and or radiotherapy) was derived from the medical record by the responsible physician from the Ghent University Center for Training and Research in Anatomical Sciences (CETRAS). Together with age, gender, sampling date and the post-mortem interval (PMI, defined as the difference between recorded time of death and sampling time), this clinical information was recorded and linked to the samples using a pseudonym (PM01, …). After sampling, subjects were embalmed and used for surgical training and educational purposes as part of the normal routine at the Anatomy and Embryology unit.

The study was carried out in accordance with the Declaration of Helsinki for experiments involving humans and was approved by the ethical committee of Ghent University Hospital (2019/1789). All donors gave written informed consent for body donation prior to death.

### Sampling, epithelial tissue isolation and DNA extraction

Small 1-2 cm² tissue biopsies were taken by experienced anatomists from 3 skin locations (upper eyelid, nose bridge and gluteal region) and 1 oral location (inner buccal region) upon arrival of the donor at the unit. Samples obtained from donors PM03-PM012 were stored at - 20°C before epithelial isolation.

After manual removal of the subcutaneous/submucosal fat, the epithelial layer was isolated using an enzymatic method, which included 4 hours incubation in a 2.5 mg/mL Dispase II solution in PBS at 37°C on a shaker plate. The epithelium was then carefully peeled away from the thicker dermis layer and circular punch biopsies (diameter 5 mm) were obtained from this thin epithelial layer (for technical reasons this procedure was reversed for PM03-PM11: first punch biopsies, then isolation). For 3 samples (i.e., 1E, 4N and 14N) larger tissue biopsies were obtained from remnant tissues. As the total biopsy surface is hard to determine for these samples, they were excluded from clone size inference.

DNA was extracted from the resulting epithelial tissue discs using the QIAamp DNA Micro Kit. Briefly, 100 µl of lysis buffer was added to each 5mm punch biopsy and subjected to lysis using the TissueLyser II (Qiagen), at speed of 30 Hz for 90 seconds. DNA was isolated according to the manufacturer’s protocol and DNA concentrations and quality were measured using NanoDrop 1000 (Thermo Fisher Scientific). DNA integrity was determined by agarose gel electrophoresis, with Hind III Digested lamda DNA (N3012S, NEB) loaded as a molecular marker.

### Deep targeted sequencing

A gene panel of 153 genes was developed for this study: 101 driver genes, 20 housekeeping genes and 32 immune genes (Table S1). The 101 driver genes included 76 genes used in an earlier study on healthy skin^4^, completed with 25 genes from Cosmic Cancer Gene Census^36^ (v91, tier 1, somatic mutations) that have been related to skin or head and neck cancer. The 20 housekeeping genes were randomly selected from a list of 3,804 previously reported genes that shown uniform expression across a panel of tissues^37^. Only genes with no previous association with cancer, as curated manually from literature, were withheld. The 32 immune genes belong to MHC pathways or have been associated with immune evasion by previous studies^31,38–42^ (Table S1). Custom baits, targeting the coding region of the gene panel, were designed using the Agilent SureDesign software on the most recent human genome build (hg38). The total size of the target regions was 566 kbp. Deep (1000x) targeted paired-end DNA nanoball sequencing (DNBseq)^43^ was performed by BGI genomics using the exome capture methodology.

### Genome alignment and somatic mutation calling

Paired-end reads (100 bp read length) were aligned to the GRCh38.p13 human reference genome using BWA-mem2^44^, BAM files from 2 different sequencing runs (lanes) were merged using Samtools^45^ and PCR duplicates were removed using Picard’s MarkDuplicates (http://broadinstitute.github.io/picard/). Somatic mutations were called using the *ShearwaterML* algorithm, available in the *deepSNV* R Package^4^. *ShearwaterML* uses a position-specific maximum-likelihood approach to call low frequency mutations from deep targeted sequencing data by comparing each sample to a set of reference samples from the same individual. The approach was applied to each sample in this study, using samples from other locations from the same donor as a reference set. Only bases with a Phred quality score >30 were analyzed to minimize the impact of sequencing errors. Mutations were detected at P_adj_<0.05 and annotated using ANNOVAR^46^. All somatic mutations located within 10 base pairs from each other were flagged for manual curation using Integrative Genomics Viewer (IGV)^47^ to check whether they occurred on the same read. If applicable, they were reannotated as DNVs, multinucleotide substitutions or deletions.

### dN/dS calculations

The ratio of non-synonymous to synonymous mutations per site (i.e., dN/dS) was calculated globally for all genes or groups of genes (driver genes, immune genes, housekeeping genes, as described higher) and individually for each gene as described previously^48^:

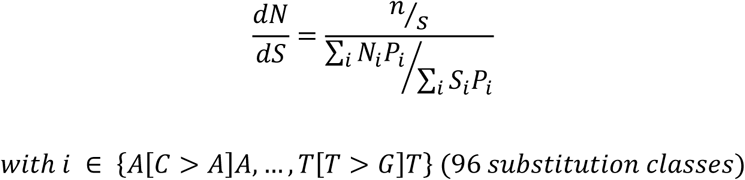

Where n is defined as the number of observed non-synonymous mutations (across all analysed skin or oral samples), s as the number of observed synonymous mutations, Ni and Si as the number of non-synonymous and synonymous sites with class i substitutions (trinucleotide substitution types) and Pi as the probability of substitution class i. Probabilities were derived from the frequencies of the 96 trinucleotide substitution classes in skin melanoma (SKCM; skin samples) and head and neck squamous skin cancer data (HNSC; oral samples), available from The Cancer Genome Atlas (TCGA, https://portal.gdc.cancer.gov/). A one-sided binomial test was used to check whether dN/dS ratios were significantly higher than 1. Genes were considered under positive selection when adjusted P (P_adj_) values (Benjamini Hochberg method) were lower than 0.1.

A related (sub)metric was calculated for missense and nonsense mutations separately by replacing the observed (n) and expected (N) number of non-synonymous mutations by the number of missense (dMiss/dS) or nonsense (dNons/dS) mutations in the formula above.

### PolyPhen-2 scores

PolyPhen-2 (PP2) scores were calculated for each mutation using ANNOVAR^46^. For each gene (or group of genes), the scores were then compared to the expected PP2 scores, calculated from 100,000 random point mutations sampled in the same gene, with prior probabilities given by the TCGA SKCM (skin samples) or HNSC (oral samples) datasets using a one-sided Wilcoxon rank sum test.

### Clone size inference

Mutational clone sizes (surface area) were estimated from the VAF using the following formula:

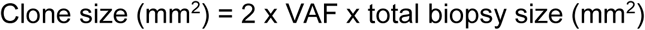

With the total biopsy size = 19.63 mm^2^ for 5 mm diameter punch biopsies.

### Healthy skin and SCC somatic mutation data

Healthy skin^4^ and SCC^27^ mutation data were downloaded from the supplementary tables provided by the authors of the respective studies. Variants were reannotated and PP2 scores added to both datasets using ANNOVAR. Genomic coordinates were converted from hg19 to hg38 using UCSC’s liftOver^49^ if required.

### Mutational signatures

The contribution of each known COSMIC SBS mutational signature to our dataset was estimated using the R Bioconductor package *MutSignatures* (function *resolveMutSignatures*)^50^.

### SiMSen sequencing

To detect and quantify 2 UV-specific hot spot mutations (Table S1), we applied SiMSen-Seq (Simple, Multiplexed, PCR-based barcoding of DNA for Sensitive mutation detection using Sequencing) as described previously^24^. Sequencing was performed on an Illumina MiniSeq instrument in 150 bp single-end mode. Raw FastQ files were subsequently processed using a modified version of Debarcer^51^. For the *RPL13A* (chr19: 49487384 - 49487466) and *DPH3* (chr3: 16264961 - 16265030) promoter amplicons, sequence reads containing the barcode were grouped into barcode families. Barcode families with at least 10 reads, where all of the reads were identical (or ≥ 90% for families with >20 reads), were required to compute consensus reads.

## Code and data availability

The targeted sequencing data that support the results reported in this study will be available at the European Genome–Phenome Archive (EGA; https://ega-archive.org), which is hosted by the European Bioinformatics Institute (EBI) and the Centre for Genomic Regulation (CRG). Due to ethical and legal reasons, the data is deposited under controlled access. Data use conditions attached to this EGA dataset limit its use to approved users at a specific institution for a specific a health/medical/biomedical project and dictate that useful results should be made available to the wider scientific community. Access requests should be addressed to Jimmy Van den Eynden.

Code and downstream data used to produce the results described in this manuscript are available on GitHub at https://github.com/CCGGlab/mutClon.

## Data processing and statistical analysis

The R statistical package (v4.0) was used for all data processing and statistical analysis. Details on the statistical tests used in this study are reported in the respective sections. *P* values less than 0.05 were considered significant for individual tests. For multiple comparisons, false discovery rate (FDR) corrections were performed using the Benjamini-Hochberg method.

## Acknowledgements

This work was supported by the Ghent University Special Research Fund Starting Grant (JVdE; BOF.STG.2019.0073.01), Kom op tegen Kanker (Stand up to Cancer), the Flemish cancer society (TL; STI.VLK.2022.0002.01) and Barncancer (JTS; TJ2021-0068). We thank all collaborators from the human Anatomy and Embryology unit at Ghent University for their daily technical and administrative support. Lastly, our deepest gratitude and thoughts are reserved for the donors and their families!

## Author contributions

J.V.d.E designed the study. J.V.d.E. and T.L. performed the bioinformatics analysis and drafted the manuscript. W.W. was responsible for donor medical record screening and inclusion and supervised donor sampling. J.V.d.V. developed the donor sampling method. A.V. and E.B. developed and performed the epithelial isolation procedure. J.S. was responsible for DNA extraction, concentration measurements and integrity checks. E.L. and K.E. performed the SiMSen sequencing. All authors read and approved the manuscript.

## Additional information

The author(s) declare no competing interests.

## Supplementary figures and tables

**Fig. S1 Detecting mutational clonality in post-mortem epithelial tissues derived from whole-body donors**.

**(a)** A 4-step methodology to detect mutational clonality. **(b)** DNA integrity gel electrophoresis image for DNA extracted from epidermal eyelid samples from 10 different subjects (indicated by circles) and with a broad range of PMIs. Donors from which samples were sequenced in this study (PM01, PM02) are indicated and labelled in red. The slightly better DNA integrity of samples obtained from these 2 subjects is likely related to the fact that these samples were obtained from fresh tissues, while the other 8 samples were obtained from frozen tissues.

**Fig. S2 Overview of all identified somatic mutations in this study**.

Somatic mutations were called from deep (1000x) targeted sequencing data using Shearwater ML. Heatmap shows all mutated genes with indication of the number of identified mutations per sample. Genes are stratified according to their type (driver genes, housekeeping genes and immune genes as indicated by the grey bar on the right) and ranked from top to bottom by mutation frequency.

**Fig. S3 Positive selection signals in driver mutations in oral and skin epithelial tissues**.

**(a)** Barplots showing dN/dS (upper panel) or dNons/dS (lower panel) for 7 genes as indicated.

**(b)** Plots showing expected (violin plot, based on simulations) and observed (scatter plots) PP2 values for 7 genes as indicated. Median values indicated by horizontal lines. For both panels, the results from this study are compared with results obtained from another study on healthy skin *(Martincorena et al. 2015*, targeted sequencing 74 genes) and with squamous cell skin cancer (SCC, whole exome sequencing) data as indicated. *, (unadjusted) *P* < 0.05; **, *P* < 0.01; ***, *P* < 0.001; NS, non-significant.

**Fig. S4 Clonal alterations in skin and oral epithelia**.

Visualization of somatic mutation-driven clones in 0.25 cm^2^ healthy skin/oral epithelium for both study donors as indicated. Clones driven by genes for which positive selection signals were found in both skin and oral epithelia are coloured as indicated in bottom legend. Other clones are uncoloured. Clones were positioned randomly with clone sizes and frequency based on somatic mutation data.

**Table S1 Main mutation data used in this study**.

Main data that underly the results reported in this manuscript, as indicated by tabnames: gene set that was used for the development of the targeted gene panel, with indication of gene type (driver, immune or housekeeping gene) and reference; mutation annotation format (maf) file; SimSen sequencing results; positive selection analysis in skin/oral epithelium with indication of dN/dS, (median) PP2 and related values.

