## Supplementary figures and images for "A clinically annotated post-mortem approach to study multi-organ somatic mutational clonality in histologically healthy tissues"

### Supplementary figures

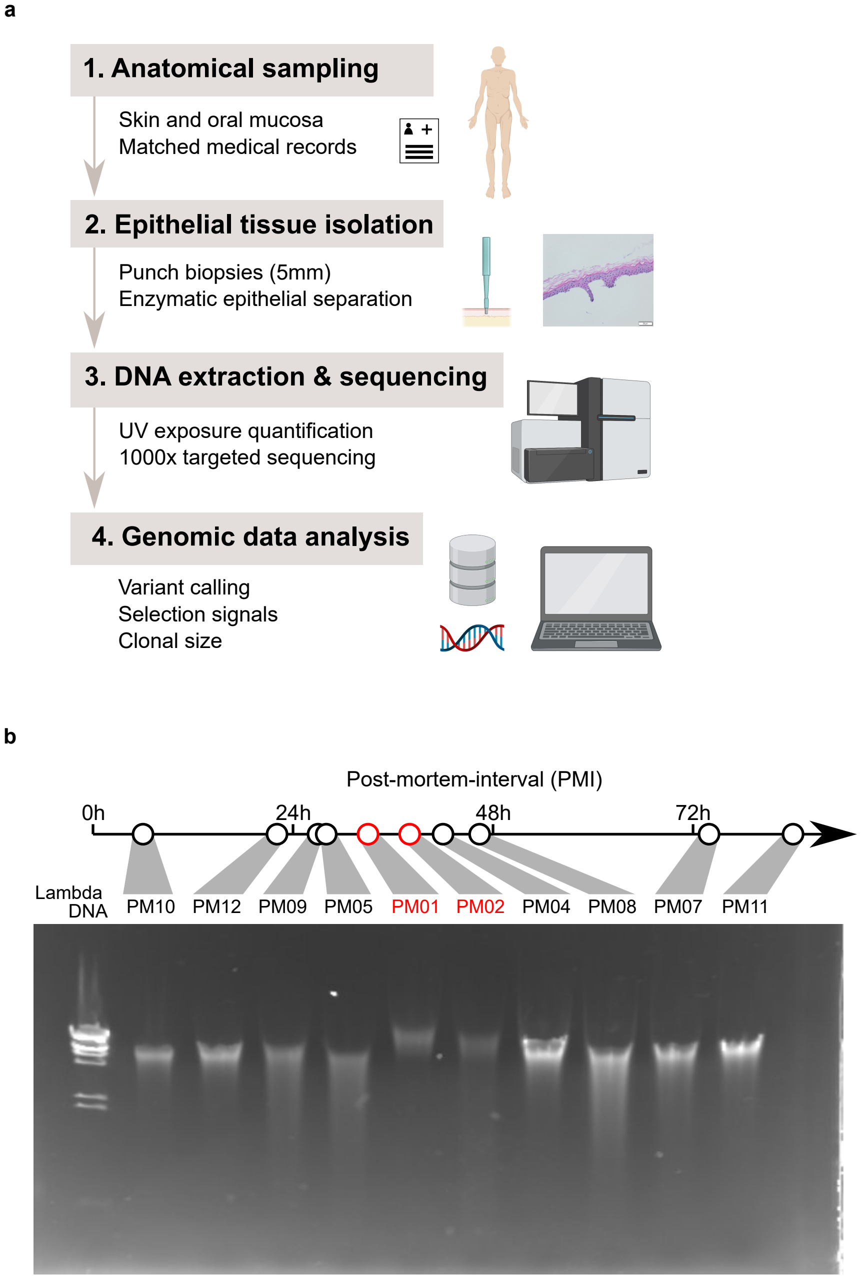


Suppl. figure 1

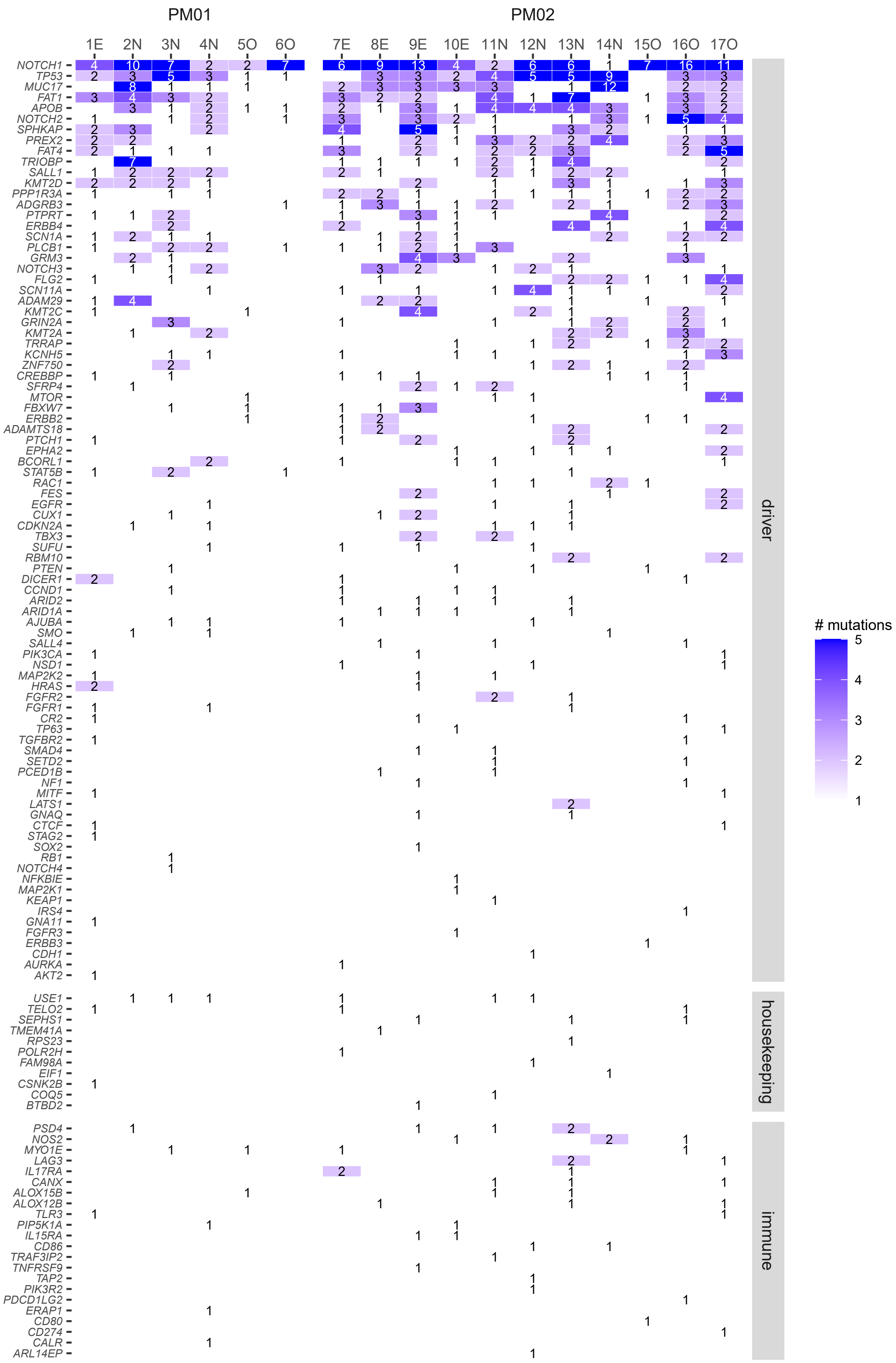


Suppl. figure 2

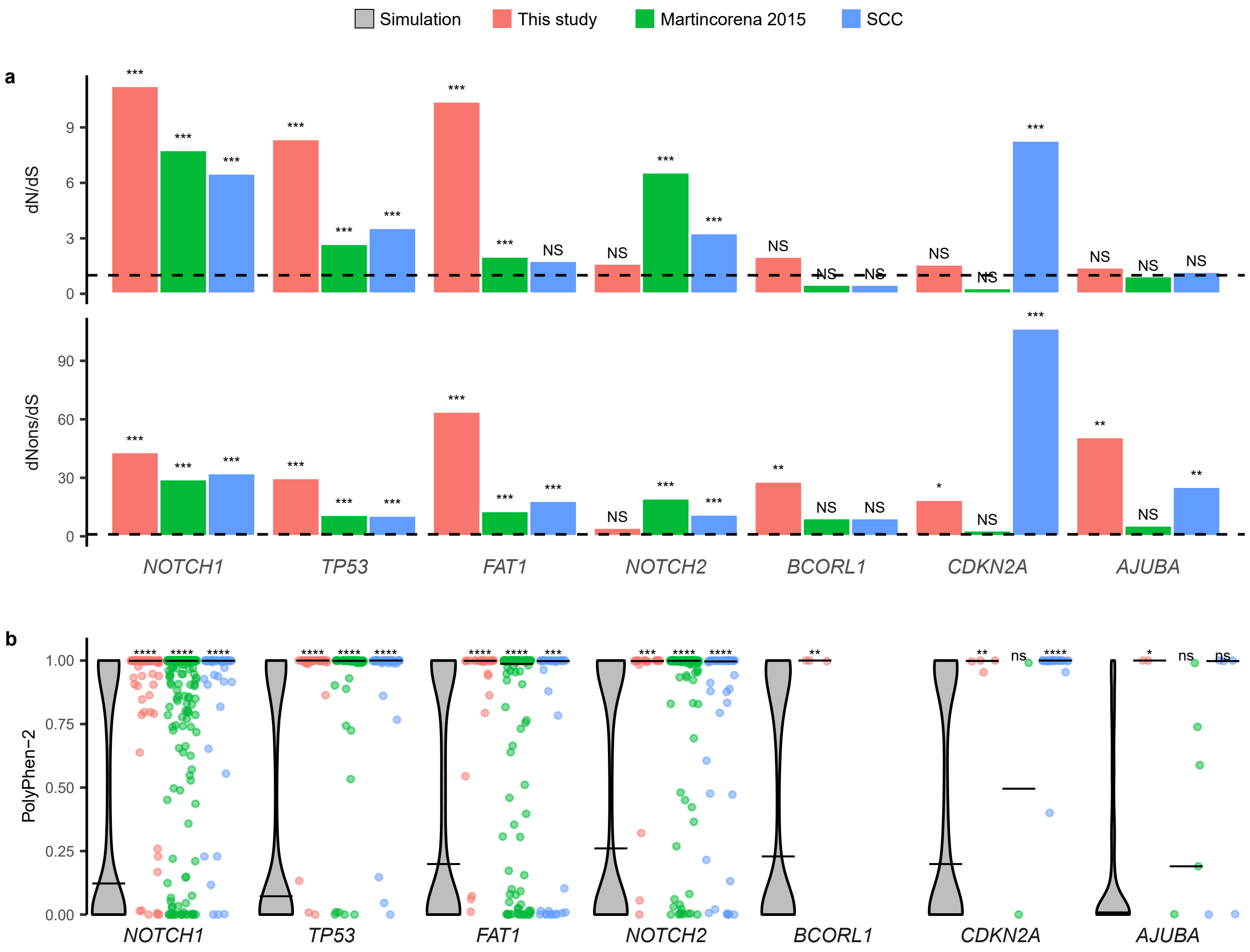


Suppl. figure 3

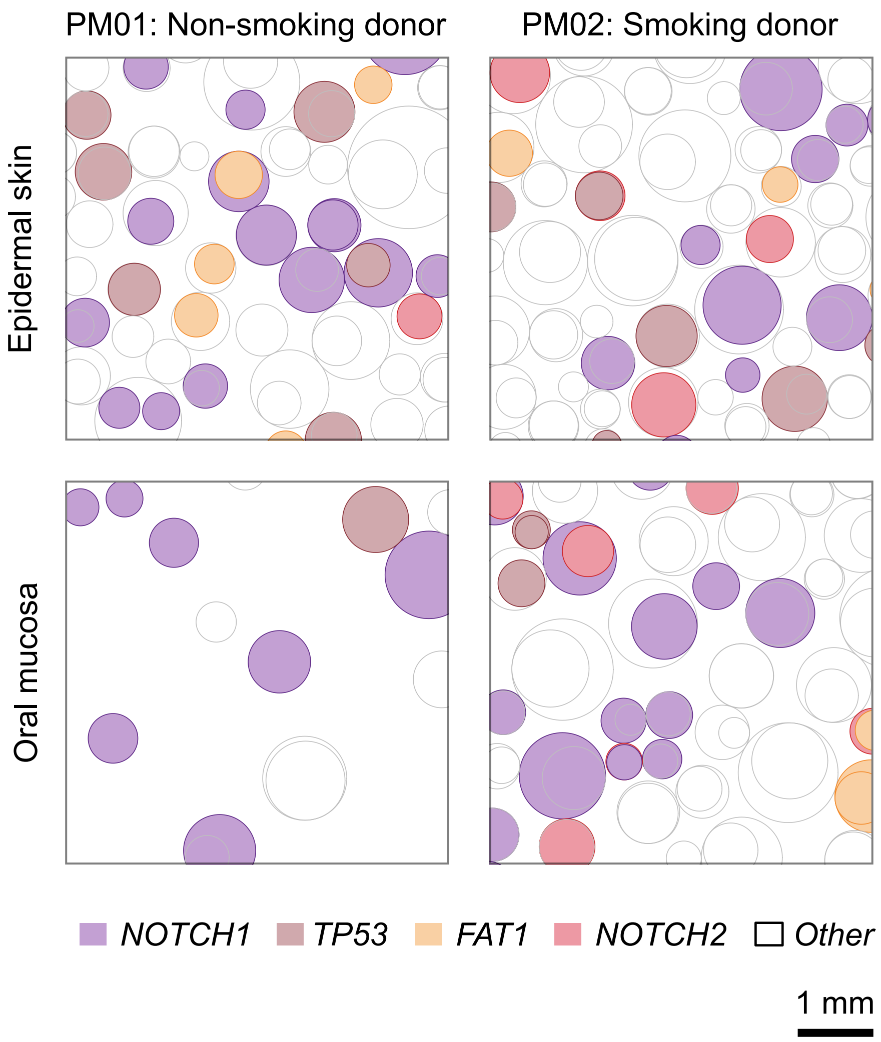


Suppl. figure 4
